# Scalable *in vivo* cardiac functional genomics with compressed AAV-Perturb-seq reveals a common mitochondrial response to perturbation

**DOI:** 10.64898/2026.06.01.729445

**Authors:** Ivan A. Kuznetsov, Kristina Li, Yijun Yang, Wencao Zhao, Wei Zhou, Wenkai Zhu, Jialiu Liang, Jian Li, Jonathan J. Edwards, Zoltan Arany

## Abstract

Efficient identification of new targets to treat human disease requires a scalable way to link genotype to phenotype directly in the target organ. Pooled CRISPR screening with single-cell RNA sequencing as a readout (Perturb-seq) has emerged as a method for functional genomics but is typically applied *in vitro* and is limited in scale. Here, we combine *in vivo* Perturb-seq via adeno-associated virus (AAV)-mediated delivery with a statistical framework allowing for signal deconvolution after multiple random perturbations per cell (compressed Perturb-seq), to develop *in vivo* compressed AAV-Perturb-seq, a scalable way to perform high-throughput, cell-autonomous functional genomics in a desired target organ. We apply this approach to study the effect of 585 gene knockouts on the cardiomyocyte transcriptome. We identify that alterations in the mitochondrial transcriptome are a common response to genetic perturbation, a finding independently validated across species and perturbation modalities. We identify very few synergistic perturbations, despite observing frequent combinatorial effects. Broadly, our work establishes a platform to facilitate *in vivo* functional genomics in a target organ with direct applicability to identifying therapeutic targets for the treatment of human disease.

## INTRODUCTION

Scalable causal linkage of cellular genetic perturbation to observed phenotype would allow for the systematic identification of new therapeutic targets. Towards this goal, pooled CRISPR screening with single-cell or single-nucleus RNA sequencing as a readout (Perturb-seq) has made it possible to study genotype-transcriptome linkage at scale and been applied to dissecting cellular regulatory circuits, studying the innate immune system, identifying autism risk genes, and exploring the phenotypic landscape of 22q11.2 deletion syndrome.^1–6^ However, large-scale conventional Perturb-seq remains prohibitively expensive and is typically implemented either *in vitro* or *ex vivo* thereby losing the critical, context-specific effects inherent to the *in vivo* environment.^7,8^ Here, we demonstrate the feasibility of circumventing these technical issues by combining adeno-associated virus (AAV)-mediated delivery of large-scale Perturb-seq libraries with a statistical framework allowing for signal deconvolution after multiple random perturbations per cell.

To do so, we build on experimental methods initially introduced by Santinha et al. with *in vivo* AAV-Perturb-seq^1^ and computational methods introduced by Yao et al. with compressed Perturb-seq/FR-Perturb.^8^ As proof of principle, we conduct *in vivo* Perturb-seq of 585 gene knockouts in the adult murine heart. Specifically, we utilize MyoAAV to deliver a sgRNA library and cTnT-driven Cre and a GFP-KASH fusion construct at low multiplicity of infection to LSL-Cas9 mice. Fluorescence-activated cell sorting (FACS) enrichment of GFP positive nuclei combined with direct sgRNA capture via ECCITE-Seq and 10X Genomics single-nuclei RNA sequencing allows for recovery of paired sgRNA-transcriptome combinations. To increase the number of genetic perturbations that could reliably be analyzed in a single experiment, we leverage the insight that single-gene perturbations tend to only affect a small number of co-regulated gene modules. Using sparsity-promoting algorithms, this allows for the deconvolution of individual perturbation effects from cells with multiple different perturbations per cell.

## RESULTS

### MyoAAV-mediated delivery allows for cardiac in vivo Perturb-seq

Recently described MyoAAV capsids^9^ allow for efficient cardiac delivery of genetic cargoes after systemic injection. To pilot this approach, we prepared AAV transfer plasmids expressing cTNT-driven Cre and GFP fused to a nuclear-localizing KASH domain (U6::sgRNA-cTnT::Cre-2A-GFP-KASH; control sgRNA) and manufactured the corresponding MyoAAV. We injected mice with escalating titers of virus, and four weeks post-infection we sacrificed mice, isolated cardiac nuclei, and conducted flow cytometry for GFP fluorescence (**Figure 1A-B; Figure S1A**). We were able to achieve between 2-23% transduction of all isolated nuclei (not exclusively cardiomyocytes; presumably ∼50% of cardiomyocytes). Next, we investigated our ability to induce indels in cardiomyocytes *in vivo*. Using medium-titer MyoAAV (MOI of 0.2-0.4), we systemically delivered to adult mice AAVs carrying sgRNAs targeting *Pank1*. Four weeks post infection we sacrificed mice, isolated cardiac gDNA, and analyzed for indel percentage. We reliably detected indels in about 13-24% of all (not just cardiomyocyte) resultant reads, demonstrating the feasibility of efficient gene knockout *in vivo* (**Figure S1B**).

**Figure 1.**
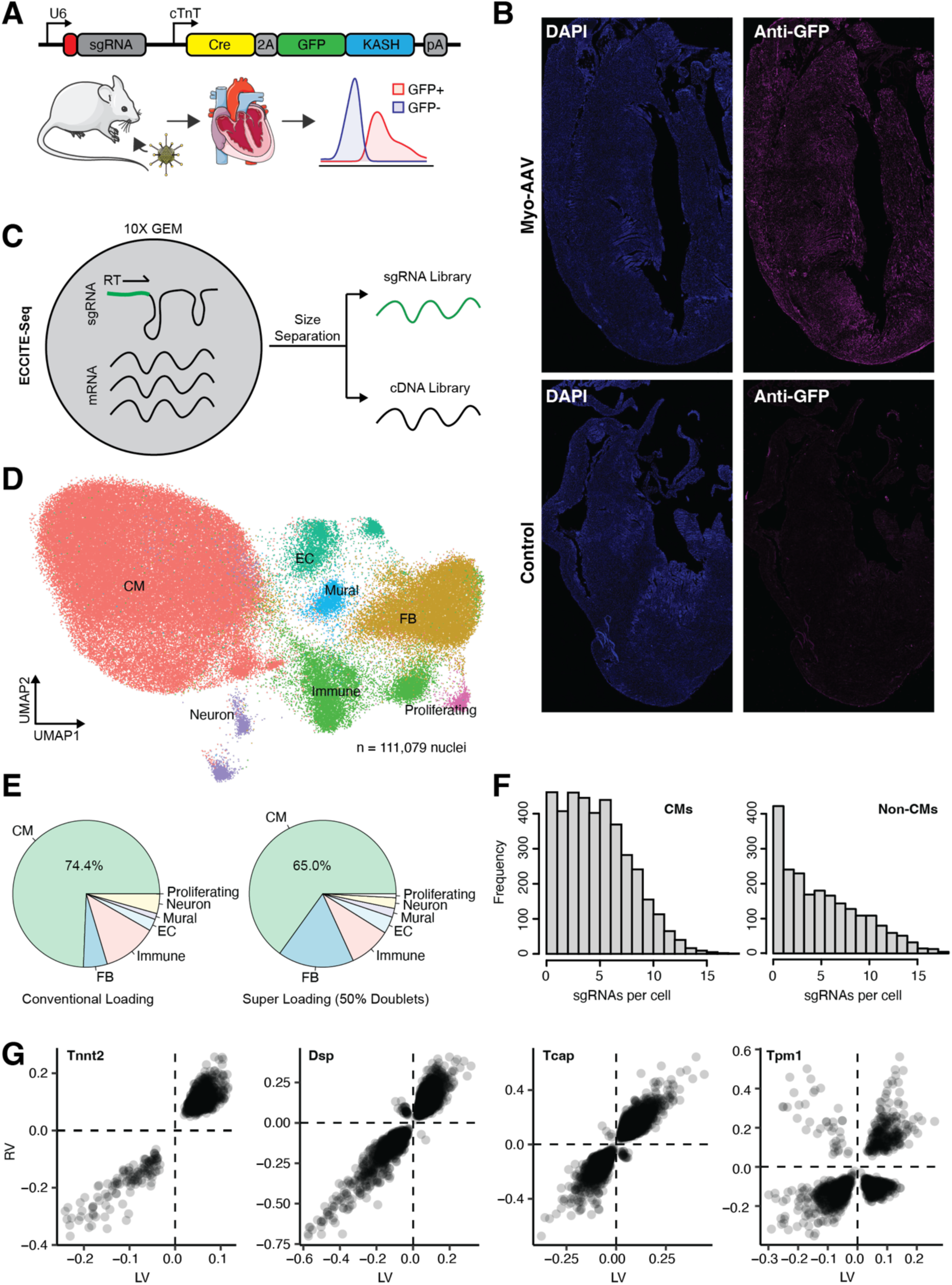
MyoAAV-mediated delivery allows for cardiac in vivo Perturb-seq. **(A)** AAV transfer plasmid design and experimental schematic. The transfer plasmid contains a U6 promoter driving transcription of an sgRNA and contains a cTnT promoter driving the transcription of both Cre and KASH-GFP. The transfer plasmid is packaged into Myo-AAV and then systemically injected into mice. After 4 weeks the hearts are harvested and processed to isolate nuclei. FACs is used to enrich for GFP positive nuclei. **(B)** Myo-AAV (5e12 vector genomes) efficiently delivers DNA to the murine heart, as seen by anti-GFP immunofluorescence. **(C)** Experimental schematic for ECCITE-seq. Briefly, an additional scaffold tracrRNA primer is added into the reverse transcription mix during the cDNA generation step to also generate a sgRNA library. The sgRNA library can then be separated from the cDNA library via size selection. **(D)** UMAP of nuclei with assigned cell type**.(E)** Proportion of each cell-type after conventional loading (targeting recovery of 5000 nuclei) versus super loading (targeting recovery of 100000 nuclei). **(F)** Number of sgRNAs assigned to each cell for cardiomyocytes versus non-cardiomyocytes. **(G)** Concordance of perturbation effects between the RV and LV for the given perturbed gene. Each point represents a downstream perturbed gene. Only perturbations which were significant are shown (q < 0.05).

Next, we piloted our ability to deliver a sgRNA library to cardiomyocytes *in vivo* and to read out sgRNA-transcriptome pairs from single-nuclei. We utilized the direct 5’ capture sequencing approach introduced with ECCITE-seq (**Figure 1C**).^10^ We cloned a library of sgRNAs targeting 5 known cardiomyopathy genes (3 sgRNAs per gene target; **Data S1**), as well as sgRNAs targeting 6 safe harbor loci, into our transfer plasmid. We then generated a pooled library of MyoAAV which we injected into mice at high MOI (MOI ∼5). 4 weeks post-transduction we harvested the murine right ventricle (RV) and left ventricle (LV), isolated nuclei, and then enriched for GFP+ nuclei via FACS prior to proceeding to ECCITE-seq. We tested two different loading strategies onto the 10X Chromium Controller: 1) conventional loading, targeting a recovery of 5,000 nuclei per lane; 2) super loading, targeting a recovery of 100,000 nuclei per lane.

Pooled across all lanes we recovered 7 different cell populations (**Figure 1D**). Approximately 75% of recovered nuclei were assigned to cardiomyocytes with conventional loading as opposed to 65% with super loading (presumably due to increased doublet rate; **Figure 1E**). We were able to assign at least one sgRNA to most cardiomyocyte nuclei and assigned an average of 5.22 sgRNAs per nuclei (**Figure 1F**). We did note some assignment of sgRNAs to non-cardiomyocyte nuclei. It was unclear if this represents contamination by ambient sgRNAs from lysed cells or if this represented mis-sorting of nuclei from non-cardiomyocyte transduced by MyoAAV. We focused our analysis on the conventional loading approach due to the cleaner enrichment of cardiomyocyte nuclei. We separately analyzed perturbation effects for the RV and LV via the FR-Perturb algorithm (**Figure S1C**) and found excellent concordance between the two across 4 of the perturbed genes (**Figure 1G**). We were underpowered to determine perturbation effects in the RV after *Myh6* perturbation due to low nuclei recovery.

Notably, we found that phenotypically the isolated murine hearts in this pilot experiment were grossly abnormal, demonstrating dilation of all 4 chambers (**Figure S1D**), presumably a consequence of wide-spread knockout of known cardiomyopathy genes. Nevertheless, this did suggest that it was important to use a lower MOI to avoid organ-level dysfunction. We were therefore faced with the challenge that we needed to keep the fraction of transduced cardiomyocytes (i.e. MOI) relatively low, while also isolating nuclei with multiple sgRNAs. We reasoned that by specifically enriching for GFP^High^ nuclei we could thereby bias our resultant nuclear suspension for nuclei transduced by more than one AAV.

### Compressed Perturb-seq identifies multifaceted in situ transcriptional perturbations in murine cardiomyocytes

We next proceeded to a full screen. We identified 585 genes of interest and generated a sgRNA library containing 4 sgRNAs per gene, as well as 500 non-targeting sgRNAs and 500 sgRNAs targeting safe harbor loci (**Data S2**). We conducted an empirical power analysis to determine the required number of perturbed nuclei (∼5,300-8,000; see **Methods**). We packaged the library into a MyoAAV capsid and injected into mice (n = 4) at a MOI of 0.2. As above, 4 weeks post-transduction we proceeded to flow sorting, this time only GFP^High^ nuclei, before conducting ECCITE-seq.

After sequencing, we determined the nuclear transcriptome using the conventional 10X CellRanger pipeline. Based on prior work demonstrating the sensitivity of Perturb-seq to sgRNA assignment, we empirically determined the optimal strategy for assigning sgRNAs to specific nuclei (see **Methods**). Once we had achieved paired sgRNA-transcriptome mappings for each nucleus (**Figure S2A-C**), we extracted perturbation effects via the FR-Perturb workflow (see **Methods**). To increase power, we conducted our analysis at the gene level, rather than at the sgRNA level. After quality control, we analyzed the transcriptome of 8663 nuclei. On average, each non-control gene was perturbed in ∼70 nuclei with ∼66% of genes being perturbed in at least 25 nuclei (**Figure 2A**). We recovered a mean of 4.63 perturbed genes per nuclei with 31.2% of nuclei contained 3 or more perturbed genes. On average, each gene in the transcriptome was affected by ∼7 perturbations, with only 0.6% of genes being unaffected by any perturbation. sgRNA assignment to non-cardiomyocytes correlated strongly with abundance of that sgRNA in cardiomyocytes, suggesting detection of ambient sgRNA (**Figure S2D**). We noted that genetic perturbation had a prominent effect on overall nuclear mRNA recovery (**Figure 2B**), with perturbation of *Mtx2*, *Dclk2*, and *Defb42* all decreasing mRNA recovery. We noted a trend that non-targeting, control sgRNAs demonstrated poorer recovery than safe-locus targeting, control sgRNAs, a surprising phenomenon not reported previously (**Figure S2E**). We observed enrichment and depletion of sgRNAs which varied over 2-3 orders of magnitude (normalized to input library sgRNA prevalence; **Figure S2F**). While we cannot rule out a contribution from differential cardiomyocyte survival or, for a small number of perturbations (e.g. *Pkm*), cell-cycle re-entry, the magnitude and distribution of the variation is most consistent with predominantly technical sources, i.e. PCR artifacts, sgRNA assignment error, and protocol-level systematic effects. The distribution of sgRNAs per nucleus was strongly right-skewed, with a minority of “super-transducer” nuclei containing ≥25 assigned sgRNAs (**Figure S3A**). Because apparent transduction load could confound perturbation-effect estimation, we characterized its transcriptomic correlates. Apparent sgRNA load correlated positively with total RNA recovered per nucleus and inversely with mitochondrial RNA content (**Figure S3B**), indicating guide and transcript capture covary. A transduction-signature score from the 406 genes enriched in ≥25 sgRNA cardiomyocyte nuclei was largely technical: UMI count and mitochondrial fraction explained 65% of its variance while sgRNA load added only ∼1% beyond these (**Figure S3C**), and a commonality analysis showed its apparent association was overwhelmingly collinear with transcript recovery rather than independent (**Figure S3D**). FR-Perturb effects for non-targeting and safe-locus control guides were near-null, showing perturbation estimates are robust to the control/normalization choice (**Figure S3E**). These analyses motivated conditioning effect estimation on transcript recovery and mitochondrial content but not sgRNA count (see **Methods**).

**Figure 2.**
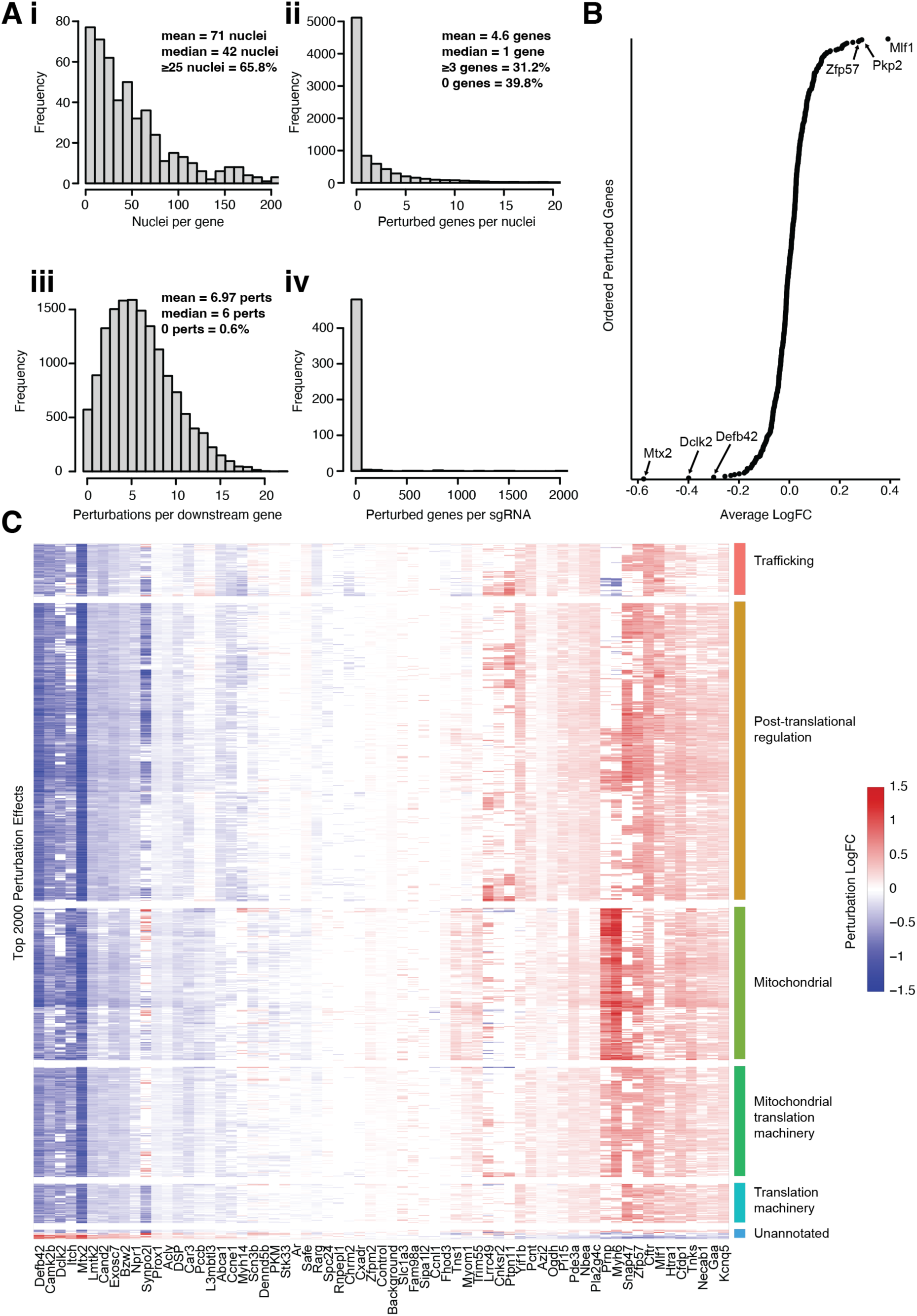
In vivo compressed AAV-Perturb Seq allows for scalable genotype-transcriptome linkage in the murine heart. **(A)** Histograms showing the distributions of the number of cells with a given gene perturbed (**i**), the number of genes perturbed per cell (**ii**), the number of downstream perturbations per perturbed gene (**iii**), and the number of perturbations per delivered sgRNA (**iv**). **(B)** Changes in overall nuclear mRNA for given perturbed gene, as normalized to nuclei with 0 sgRNA. **(C)** Top 2000 perturbation effects for given perturbed genes, organized by corresponding gene modules. Only perturbed genes with at least 100 downstream perturbations with q > 0.5 are shown. Perturbations with q < 0.5 are shaded white.

After perturbation recovery, we identified 65 genes that upon perturbation generated at least 100 significant downstream effects (as compared to nuclei with 0 sgRNA; **Figure 2C**). We clustered perturbation effects into 6 different gene modules (trafficking, post-translation modification, translational machinery, nuclear-expressed mitochondrial genes, mitochondrial translation machinery, and a small unannotated group; see **Methods; Data S3**).

### Alteration of the mitochondrial transcriptome is a common downstream result of perturbation

We found that alterations of the nuclear-synthesized mitochondrial transcriptome was a common response to perturbation, with 23 perturbations demonstrating significant effects (**Figure 3A-B**). Analysis by gene function demonstrated that although all aspects of mitochondrial homeostasis were altered by perturbation, the most affected were oxidative phosphorylation transcripts (**Figure 3C**). Specifically, 15 of the 23 perturbations which affected the mitochondrial transcriptome demonstrated significant effects onto oxidative phosphorylation transcripts. Notably, many of the 23 perturbations are not themselves mitochondrial genes, suggesting that mitochondrial transcriptome dysregulation represents a convergent downstream response to diverse genetic insults in cardiomyocytes.

**Figure 3.**
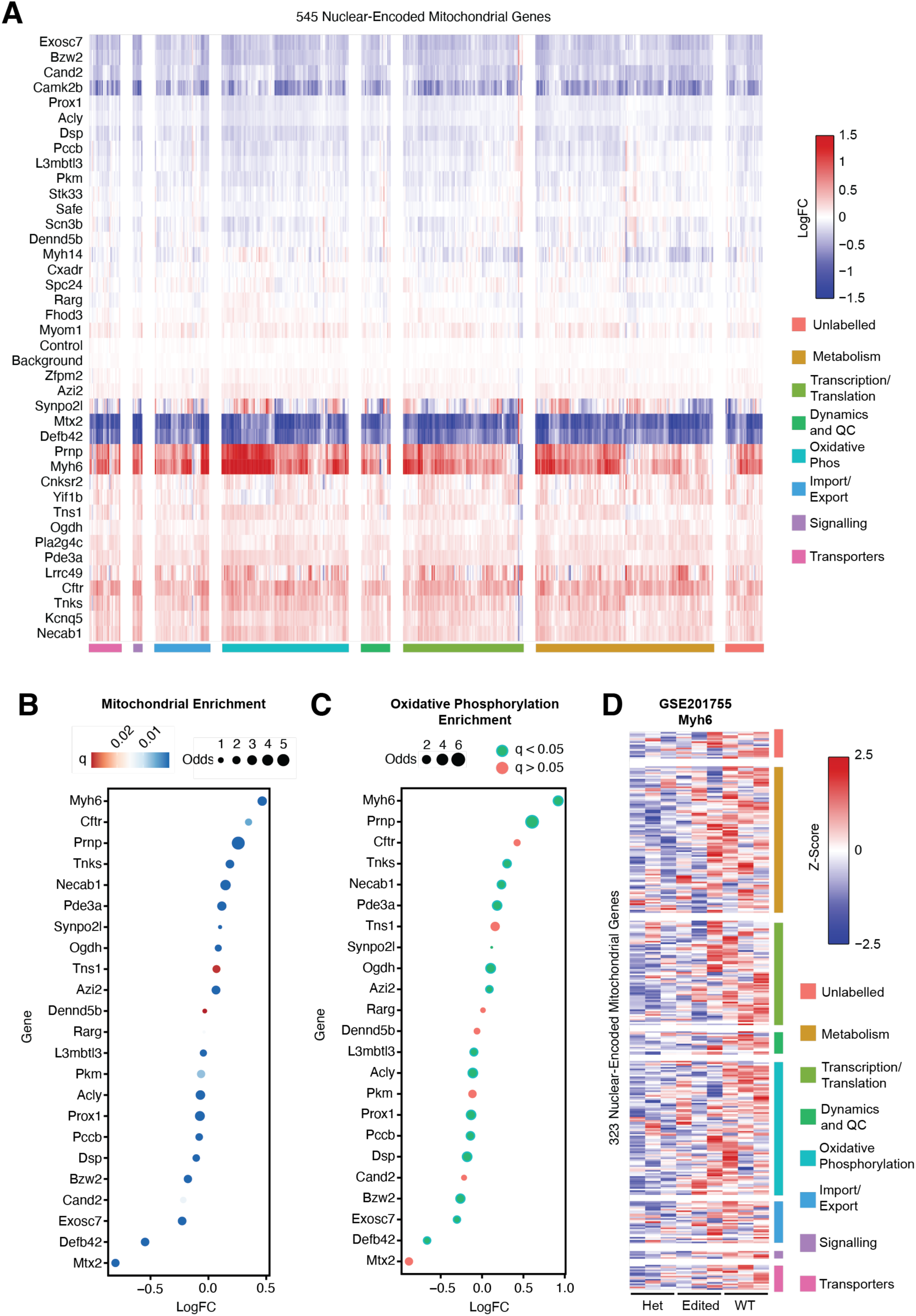
Alteration of the mitochondrial transcriptome is a common response to genetic perturbation in the murine heart. **(A)** Log fold change effects of perturbations on the mitochondrial transcriptome, grouped by gene function. Only perturbations with at least 100 downstream effects with q > 0.5 are shown. Perturbations with q < 0.5 are shaded white. **(B)** 23 perturbations significantly influenced nuclear-synthesized mitochondrial genes. Here, a perturbation was considered as having a significant effect on a downstream gene if q < 0.2. Odds ratios and p-values are calculated using Fischer’s exact test and q-values are generated using the Benjamini-Hochberg correction. **(C)** Specific effect on oxidative phosphorylation transcripts (versus all other mitochondrial transcripts) for the 23 perturbations with significant effects on the mitochondrial transcriptome. **(D)** The mitochondrial transcriptome is downregulated in a humanized hypertrophic cardiomyopathy mouse model (*Myh6^h403/+^*) in which the human MYH7 c.1208G>A (p.R403Q) pathogenic variant is knocked into the endogenous murine *Myh6* locus and is partially restored by AAV base-editing correction. Heatmap shows z-scored expression of MitoCarta^26^ genes (among the top 2,000 most perturbation-responsive genes) in wild-type, heterozygous *Myh6^h403/+^*, and AAV base-editor–treated hearts, grouped by MitoCarta functional category.

First, we cross-referenced our findings with bulk RNA-seq data from a humanized hypertrophic cardiomyopathy mouse model (*Myh6^h403/+^*) in which the human *MYH7* c.1208G>A (p.R403Q) pathogenic variant is knocked into the endogenous murine *Myh6* locus and subsequently corrected by AAV-delivered base editing (GSE201755).^11^ Re-analysis revealed that heterozygous *Myh6^h403/+^* mice exhibited broad downregulation of nuclear-encoded mitochondrial transcripts relative to wild-type, with the most pronounced effects concentrated in oxidative phosphorylation genes, closely paralleling the mitochondrial transcriptome signature we identified for *Myh6* perturbation. Notably, AAV base-editing correction partially restored this mitochondrial gene expression pattern toward wild-type levels, suggesting that the link between sarcomere dysfunction and mitochondrial transcriptome dysregulation is reversible and causally downstream of the primary mutation (**Figure 3D**).

Second, to further validate that our FR-Perturb signatures accurately reflect gene-level transcriptional changes, we compared our perturbation effect sizes against published bulk RNA-seq datasets from independent cardiac knockout models. For cardiomyocyte-specific *Dsp* conditional knockout (GSE180972^12^), FR-Perturb log-fold changes were weakly correlated with those measured in the published knockout (Pearson r = 0.18, permutation p < 0.001, n = 1041 mitochondrial genes), with oxidative phosphorylation genes showing the most pronounced downregulation (**Figure S4A–B**). Similarly, FR-Perturb signatures for *Pkm* knockdown were weakly correlated with published *Pkm2* cardiac knockout data at baseline (GSE243668^13^; Pearson r = 0.10, permutation p < 0.001; **Figure S4C–D**), with oxidative phosphorylation genes again showing small, but significant, downregulation. These results demonstrate that FR-Perturb recovers biologically coherent gene-level signatures that are reproducible in independent *in vivo* loss-of-function models.

Third, we cross-referenced our findings with Takeuchi et al., which reported a large-scale CRISPRi Perturb-seq screen in human iPSC-derived cardiomyocytes.^14^ Takeuchi et al. systematically profiled the transcriptional consequences of knocking down 1,983 transcription factors, identifying 250 co-regulated gene programs that collectively map the transcription factor regulatory landscape of the human cardiomyocyte. Of the 250 gene programs identified in that study, we found 6 programs enriched for mitochondrial genes. Of the top 10 most frequently perturbed programs, 2 were mitochondrial programs (**Figure S4E**). 36.6% of transcription factor perturbations affected at least one of these mitochondrial programs, closely paralleling our observation that 35.4% of perturbations (23/65) affected the mitochondrial transcriptome (**Figure S4F**).That this convergence on mitochondrial transcriptome dysregulation was observed independently across species (mouse and human), perturbation modalities (CRISPR knockout and CRISPRi knockdown), and perturbation gene classes (mixed functional categories and transcription factors only) suggests it reflects a fundamental property of the cardiomyocyte regulatory network.

Towards identifying the functional significance of the observed perturbation effects, we proceeded to integrate our Perturb-seq results with GWAS data using sc-linker.^15–17^ As has been previously done, we hypothesized that if a perturbed gene is important for a given trait, then disease heritability should be enriched near its downstream perturbed genes.^8^ We constructed two programs for each perturbed gene which either reflects a weight set of upregulated or downregulated downstream genes, with weights reflecting the effect size of the perturbation on that downstream gene. After filtering, we retained 123 programs for downstream heritability enrichment. We identified 7 program/trait pairs which demonstrated significance at a modest cutoff of q < 0.2 (**Figure S5A**). This analysis implicated *Myh6* (upregulated genes), *Pccb* (downregulated genes), *L3mbtl3* (downregulated genes), and *Itch* (downregulated genes) with hypertrophic cardiomyopathy (HCM) and *Itch* (downregulated genes) with a propensity for the development of atrial fibrillation. *Myh6* encodes α-myosin heavy chain, which is the dominant adult ventricular isoform in mice (functionally analogous to MYH7/β-myosin heavy chain in humans); rare *MYH6* variants have been linked to HCM in humans.^18,19^ *Itch* has previously been reported to suppress cardiac-hypertrophy via the Wnt/β-catenin signaling pathway.^20^ Mutations in *Pccb* are associated with propionic acidemia and can cause cardiomyopathy.^21^ *L3mbtl3* polymorphisms have been associated with coronary artery disease but not with cardiomyopathy specifically.^22^ Notably, on examining the downstream perturbations driving the enrichment for HCM, we found that many of them were mitochondrial genes (**Figure S5B**).

### Reconstruction of the murine RVF transcriptome using perturbation signatures

A key challenge in functional genomics is the inverse problem, i.e. inferring the set of perturbations which lead to a specific observed phenotype.^23^ Biological complexity and degeneracy make this problem generally intractable. However, occasionally finding a specific sparse solution under stringent assumptions is still useful. In this context, we use sparse to mean a minimal perturbation set which yields the phenotype of interest. Towards this, we reasoned that the transition from a healthy RV to a failing RV after pulmonary artery banding (PAB) could be modeled as a linear combination of multiple perturbations. We analyzed prior snRNA-seq data from murine RVs after PAB (n = 3 sham; n = 3 moderate RVF, n = 4 severe RVF), analyzing only genes which were differentially expressed between the two conditions.

We jointly inferred perturbations across samples using a multi-task sparse regression framework, enforcing shared sparsity while allowing sample-specific effect sizes (see **Methods**). We initially pooled perturbations into 146 potentially overlapping GO modules (requiring at least 10 genes per module; **Table S4**) and fit the severe RVF transcriptome in a lower dimensional subspace so as to increase the smoothness of our objective function. We found that a weighted combination of 42 module perturbations (representing 275 gene perturbations) demonstrated Pearson correlation of 0.627 (0.612, 0.651, 0.602, 0.643 across the 4 mouse samples) with the observed severe RVF transcriptional shift (**Figure 4A-B**). We then focused specifically on these 275 genes and refit the data using gene-level perturbations, this time more heavily enforcing sparsity. We found that a weight combination of 41 gene perturbations demonstrated 0.610 correlation (0.606, 0.636, 0.577, 0.621 across the 4 samples) with the observed severe RVF transcriptional shift (**Figure S6A-C**). The correlation is significantly greater than what would be observed based on chance alone, even without enforcing sparsity (**Figure S6D-E**).

**Figure 4.**
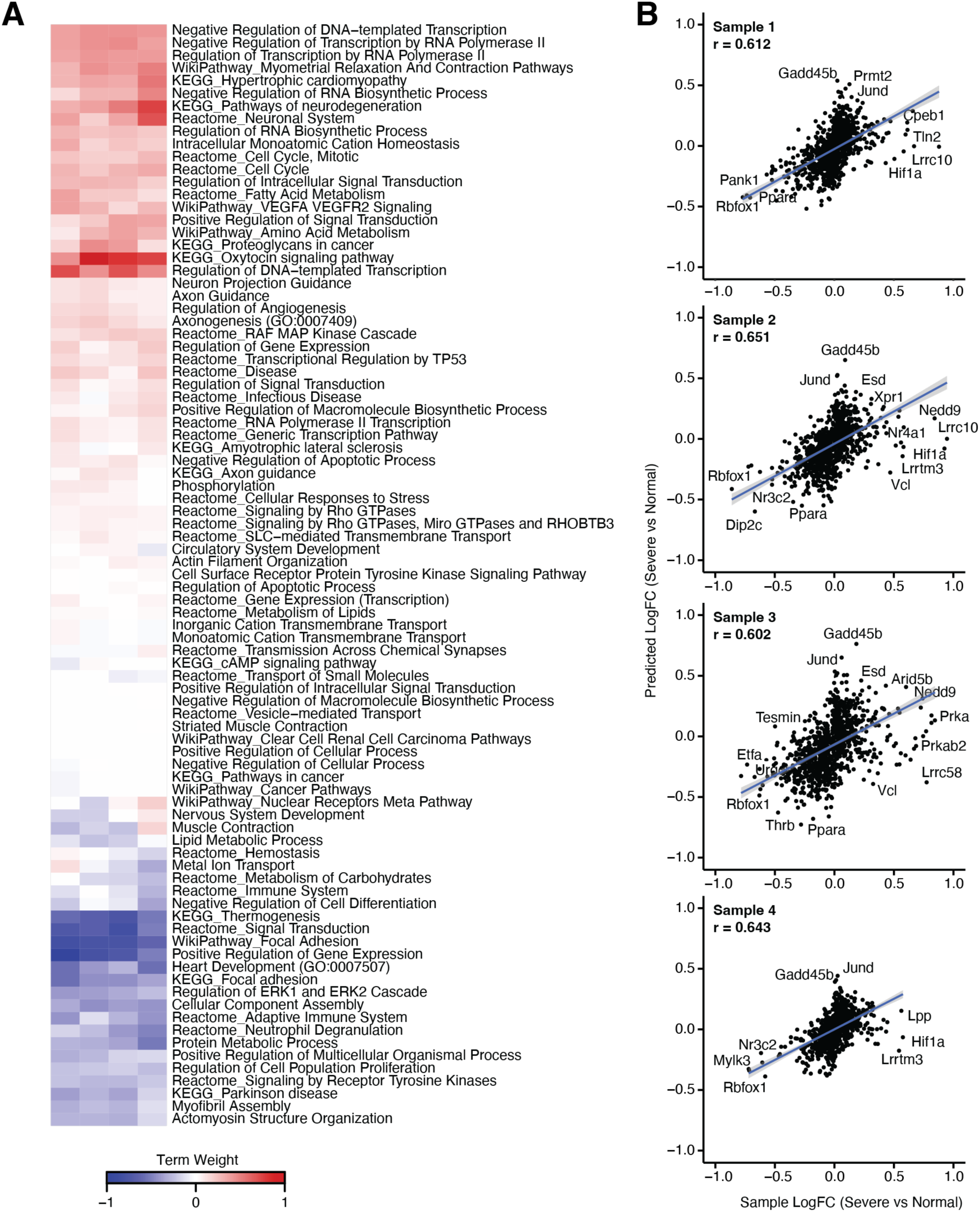
Reconstruction of the murine RVF transcriptome using gene module perturbation signatures. **(A)** Heatmap showing weights of the elastic net (alpha = 0.1) regression minimized gene module perturbation set which replicates the transcriptional signature of murine RVF. **(B)** Dot plots showing measured versus fit fold changes between healthy and failing murine RVs across 4 samples.

To assess whether the murine perturbation-derived RVF signature generalizes beyond the training data, we next asked whether a failure signature score, calculated from the fixed lasso weights, could stratify PAB mice by disease severity. We observed a gradual increase in failure score with disease severity (**Figure 5A**). These lasso weights, learned entirely from mouse PAB cardiomyocytes without any human data, could stratify independent human RV samples by disease severity: applying the original model weights to pseudobulk transcriptomes from human snRNA-seq across non-failing (NF), pressure-loaded/non-failing RVs (pRV), and RVF cardiomyocytes without any refitting yielded a significant graded increase in signature score across disease severity groups (Spearman ρ = 0.71, Kruskal-Wallis p = 0.016; **Figure 5B**). This trend was not recapitulated by permuted models in which the gene-weight assignments were shuffled (p < 0.001; **Figure 5C**), nor by models using random perturbation sets of matched size (p = 0.035; **Figure 5D**), confirming that it is the specific combination of perturbation signatures, rather than the number or magnitude of weights, that drives cross-species stratification. That a signature derived exclusively from murine *in vivo* perturbation data can stratify human RVF severity without retraining suggests a conserved transcriptional architecture underlying right heart failure across species. However, the specific identity of the selected perturbations was not uniquely required for cross-species generalization. When we refit model weights to the murine PAB data using randomly sampled 41-gene perturbation sets, generalization to the human data was not significantly worse than the original set (p = 0.109; **Figure 5E**), although the original set still outperformed ∼89% of random refits. This is consistent with the right ventricular failure transcriptome being sufficiently low-dimensional that multiple distinct perturbation combinations can approximate it with comparable fidelity (see Discussion).

**Figure 5.**
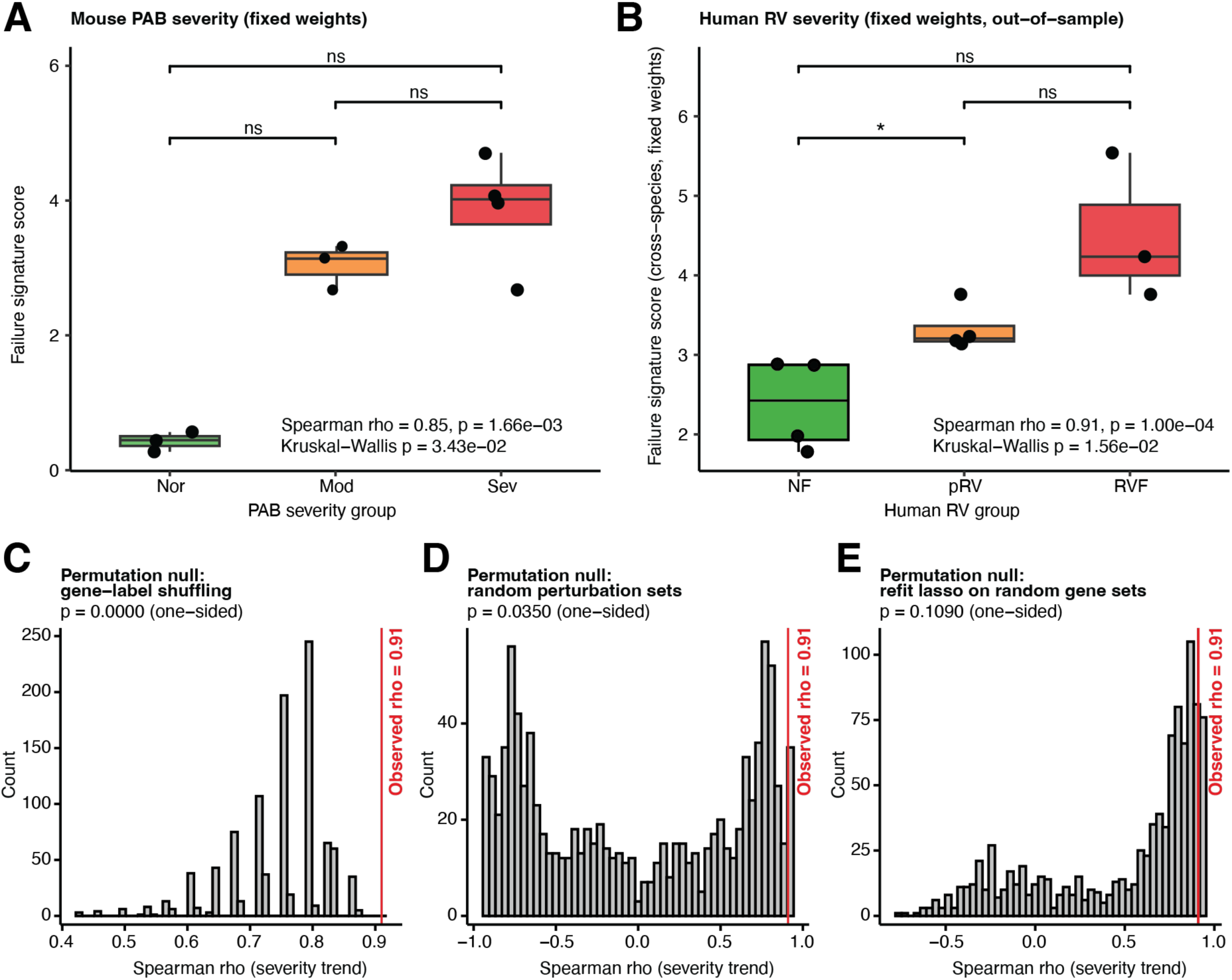
Out-of-sample cross-species validation of the murine perturbation-derived RVF signature. **(A)** Failure signature scores in mouse PAB pseudobulk samples stratified by hemodynamic severity (Normal, Moderate, Severe). Scores are computed by projecting pseudobulk cardiomyocyte transcriptomes onto the fixed lasso-derived perturbation signature. Each point is one pseudobulk sample; boxes show median ± IQR. Significance brackets indicate Wilcoxon rank-sum tests between adjacent severity groups. **(B)** The same fixed perturbation weights, applied without refitting to independent human right ventricular snRNA-seq pseudobulk samples from non-failing (NF), pressure-loaded but normal function RV (pRV), and failing RV (RVF) donors. Spearman ρ and Kruskal-Wallis p-value assess the monotonic severity trend. No human data were used in model training. **(C)** Permutation null distribution for the human severity trend (Spearman ρ) generated by shuffling gene-weight label assignments 1,000 times. Red vertical line indicates the observed ρ. One-sided permutation p-value tests whether the specific gene–weight pairings, rather than the weight magnitudes alone, drive the observed trend. **(D)** Permutation null distribution generated by replacing the 41 model perturbations with randomly sampled perturbation sets of equal size and assigning the original weight values, repeated 1,000 times. Tests whether the identity of the selected perturbations is necessary for cross-species stratification. **(E)** Permutation null distribution generated by refitting the model weights with 41 randomly selected genes to the PAB data and testing on the human data, repeated 1,000 times. Tests the uniqueness of the identity of the selected perturbations in predicting RVF severity.

### Second-order genetic interactions are common in cardiomyocytes in vivo

Leveraging the presence of multiple perturbation per cell, we sought to identify non-additive genetic interactions in the perturbed transcriptome of cardiomyocytes. We were not powered to analyze interaction effects at the gene level due to the combinatorial complexity of second-order interaction effects (170,820 perturbation pairs). Instead, using the approach proposed by Yao et al., we leveraged our prior 146 aggregated GO annotation modules.^8^ We also grouped downstream perturbed genes by their program membership. We then calculated intra-module second-order interaction effects as the deviation from summative effects in cells containing perturbations for two genes within the same module. Similarly, we calculated inter-module effects as the deviation from summative effects in cells containing perturbations for two genes from different modules. For the inter-module effect calculation, we further clustered the GO modules into 8 super-modules (28 super-module pairs) before proceeding with the analysis to improve statistical power.

We detected 29 modules with significant (q < 0.05) intra-module interactions for at least one gene program (**Figure 6A-C**). 24 of the 28 possible inter-module pairs showed significant (q < 0.05) interactions (**Figure 6D-F**). Interestingly, nearly all second-order interactions tended to be suppressive, causing the combined effect of 2 sgRNAs to be sub-linear. We observed only 2 intra-modular and 0 inter-modular synergistic effects.

**Figure 6.**
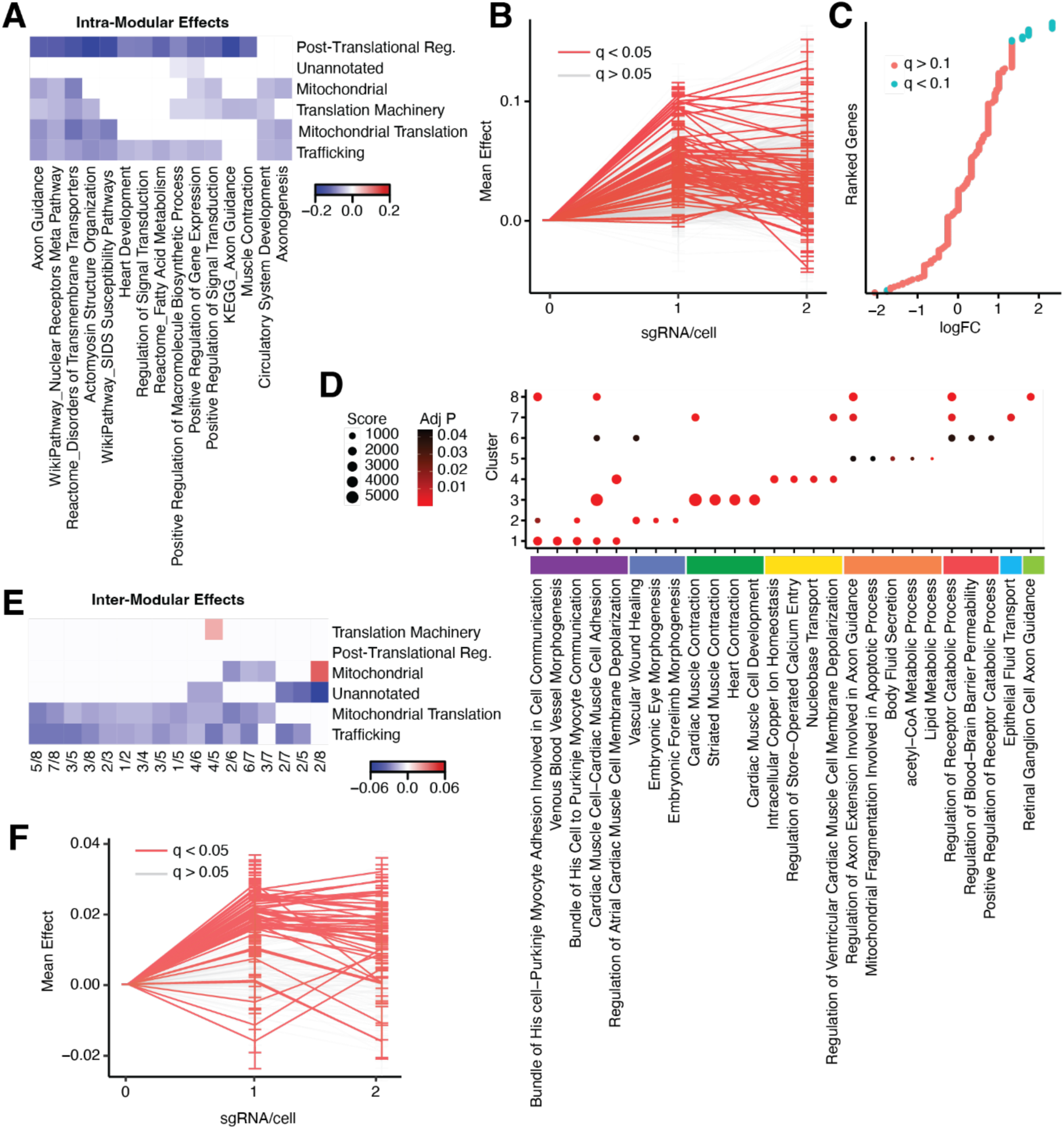
Second-order genetic interactions are common in cardiomyocytes in vivo. **(A)** Heatmap showing mean effect size of second-order intra-modular effects. Only GO terms with significant intra-modular effects (q < 0.05) for at least 2 gene modules are shown. **(B)** Visual representation of intra-modular second-order effects, showing mean effect size for 0, 1, or 2 sgRNAs and apparent deviation from linearity, i.e. 2 sgRNAs leads to double the effect of 1 sgRNA. Significant deviations from linearity are colored in red. **(C)** Certain genes are over/under-represented (colored blue if q < 0.01) in GO terms which show deviation from linearity. **(D)** Leiden-clustering of GO terms into perturbation modules yields 8 clusters with shown GO enrichments. **(E)** Heatmap showing mean effect size of second-order inter-modular effects. Only GO terms with significant intra-modular effects (q < 0.05) for at least 2 gene modules are shown. **(F)** Visual representation of inter-modular second-order effects, showing mean effect size for 0, 1, or 2 sgRNAs and apparent deviation from linearity.

## DISCUSSION

### *In Vivo* Perturb-seq as a platform for functional genomics

Despite rapid growth in the adoption of Perturb-seq approaches for functional genomics, *in vivo* applications have remained limited. This is partially due to Perturb-seq approaches relying on lentiviral delivery and genomic integration for purposes of readout. However, *in vivo* approaches rely primary on AAV-mediated delivery, which raises the challenge of reading out sgRNA presence without genomic integration. Another issue is one of scale – large scale conventional Perturb-seq studies require 100,000s of cells, which is limiting both in term of reagents and sequencing costs. Combining direct sgRNA capture (e.g. as in ECCITE-seq) with guide-pooled compressed Perturb-seq (e.g. FR-Perturb) addresses both these issues, allowing us to study ∼600 gene perturbations across 4 10X flow lanes.

*In vivo* perturbation offers several advantages that are difficult to recapitulate *in vitro*. Most importantly, it enables the study of gene function within the native cellular and tissue context, where cells experience physiologically relevant signaling, architecture, and microenvironmental cues. This is particularly critical for processes that depend on cell–cell interactions, extracellular matrix composition, or systemic factors that are absent or altered in culture. Additionally, *in vivo* approaches allow for perturbation with minimal disruption to tissue organization, avoiding artifacts introduced by dissociation, culture adaptation, or supraphysiologic delivery conditions. Together, these factors make *in vivo* Perturb-seq approaches uniquely suited to improving the fidelity and interpretability of functional genomics studies.

Several practical lessons emerged from our implementation that are relevant for future *in vivo* Perturb-seq efforts. First, we observed that high MOI delivery of a library enriched for known cardiomyopathy genes induced gross cardiac chamber dilation, indicating that widespread mosaic knockout of disease-relevant genes can produce organ-level dysfunction that confounds cell-autonomous interpretation. Maintaining a low MOI is therefore critical, but this creates a tension with compressed Perturb-seq, which benefits from multiple perturbations per cell. Our strategy of enriching for GFP^High^ nuclei via FACS partially addresses this trade-off by biasing the recovered nuclear population toward multiply-transduced cells without requiring high systemic MOI. However, we note that this enrichment strategy is not without potential bias. To assess whether this enrichment biases the recovered population, we characterized the transcriptomic correlates of apparent transduction load. Although 406 genes were elevated in heavily transduced (“super-transducer”) cardiomyocyte nuclei, this signature proved to be overwhelmingly technical rather than biological: it is largely explained by transcript recovery and mitochondrial content, with apparent sgRNA load contributing little independent variance once these are accounted for (**Figure S3C–D**). We therefore conditioned perturbation-effect estimation on these recovery covariates rather than on sgRNA count itself, which proxies the perturbation exposure and would constitute over-adjustment (see **Methods**). After this adjustment the residual transduction-associated signal is small and largely confined to the extreme super-transducer tail. Nonetheless, because GFP^High^ enrichment skews the recovered population toward more efficiently captured nuclei, we cannot fully exclude a modest sampling bias toward a recovery- or quality-distinct subset of cardiomyocytes, which warrants consideration in the interpretation of our results and in future experimental designs. As additional technical caveats, we note that sgRNA assignment to non-cardiomyocytes scaled with cardiomyocyte sgRNA abundance, most consistent with ambient contamination although true non-cardiomyocyte transduction cannot be fully excluded (**Figure S2D**). Additionally, non-targeting controls recovered more poorly than safe-locus controls, an unexpected effect that may inform control-guide choice in future compressed screens (**Figure S2E**). Overall, while FR-Perturb’s sparsity assumptions confer robustness to noise, systematic sgRNA-assignment error could still propagate into the deconvolution of perturbation effects, particularly for genes with low representation in the library.

More broadly, the compressed Perturb-seq framework relies on the assumption that individual perturbation effects are sparse, affecting only a small number of co-regulated gene modules, and that the transcriptional response to multiple simultaneous perturbations can be linearly decomposed into individual effects. Our analysis of second-order interactions provides empirical support for the approximate validity of the linearity assumption, though the predominantly sub-linear nature of observed combinatorial effects suggests that compressed deconvolution may systematically overestimate the magnitude of individual perturbation effects in multiply-perturbed cells. In biological contexts where epistatic interactions are stronger or where perturbations have large non-cell-autonomous effects (for example, in tissues with extensive paracrine signaling) the assumptions underlying FR-Perturb may be more substantially violated, and alternative deconvolution strategies may be required.

Looking forward, the platform described here is not inherently limited to the heart. The core experimental framework, i.e. AAV-mediated delivery of a sgRNA library to LSL-Cas9 mice followed by FACS enrichment and ECCITE-seq, is in principle generalizable to any organ for which an efficient AAV serotype and a tissue-specific promoter are available. Liver, skeletal muscle, and retina represent particularly tractable targets given the high transduction efficiency achievable with existing AAV capsids (e.g. AAV8 for liver, MyoAAV for skeletal muscle). Extension to the central nervous system is also feasible, as demonstrated by Santinha et al.^1^ More broadly, as engineered AAV capsids with enhanced tissue tropism and reduced immunogenicity continue to be developed, the range of organs accessible to in vivo compressed Perturb-seq will expand correspondingly. The principal remaining bottleneck is not the delivery itself but rather the cost and throughput of single-nucleus sequencing, which currently limits the number of perturbations that can be powered in a single experiment. Continued advances in combinatorial indexing, split-pool barcoding, and other ultra-high-throughput sequencing methodologies may ultimately relax this constraint.

### Mitochondrial transcriptome dysregulation as a convergent response to genetic perturbation

A common and preserved transcriptional response to perturbation in cardiomyocytes was alterations in the nuclear-synthesized mitochondrial transcriptome, with oxidative phosphorylation transcripts disproportionately affected. This finding was independently validated across orthogonal datasets. Mitochondrial transcripts were similarly altered in a humanized hypertrophic cardiomyopathy mouse model in which the human MYH7 c.1208G>A (p.R403Q) pathogenic variant was knocked into the endogenous murine *Myh6*.^11^ In a large-scale CRISPRi screen of 1,983 transcription factors in hiPSC-derived cardiomyocytes^14^ 36.6% of transcription factor perturbations affected mitochondrial gene programs, closely paralleling the 35.4% observed in our screen. That this convergence was observed across species and perturbation modalities suggests that mitochondrial transcriptome dysregulation is a fundamental feature of the cardiomyocyte stress response rather than an artifact of any single experimental system. Whether this reflects a direct regulatory coupling between diverse gene networks and mitochondrial biogenesis, or a secondary consequence of disrupted cellular energetics, remains to be determined.

### Second-order combinatorial effects are common but synergy is rare

Our experimental design to achieve multiple perturbations within a single cell was naturally suited to the analysis of second-order combinatorial effects. We found that deviations from linearity were common, but that these deviations made the system behave sub-linearly. Interestingly, nearly all detected second-order interactions were suppressive, with combined effects falling below the linear expectation. We observed only 2 intra-modular and 0 inter-modular synergistic effects. This stands in contrast to studies in other systems where both synergistic and buffering interactions are commonly observed,^24^ and suggests that the cardiomyocyte regulatory network may be particularly robust to combinatorial genetic insult. That epistatic interactions non-additively regulate cardiac phenotypes has recently been demonstrated in the context of cardiac hypertrophy,^25^ though the predominance of buffering over synergy that we observe in the transcriptome has not, to our knowledge, been previously reported.

### Reconstruction of a complex biological phenotype via *in silico* combinatorial perturbation

Conceptually, disease processes reflect a collective deviation from a baseline healthy state, that can be captured transcriptionally as an integrated function of a multitude of individual gene perturbations. Unraveling this integrated disease function has implications that could be leveraged for therapeutic development. Several factors, however, make this generally impractical. For example, uniqueness of any individual genetic perturbation cannot be assumed, and instead several distinct genetic perturbation sets could converge to an equivalent biological phenotype. Perhaps an even greater barrier is that we have very little *a priori* knowledge regarding the landscape of this integrated function. Here, we show that we can fit the murine RV cardiomyocyte transcriptomic response to PAB under the very strict assumptions that 1) the response to gene knockdown is simply a scaler weighting of the response to knockout, 2) the response to gene upregulation is the inverse of the response to downregulation, and that 3) the response to a multi-gene perturbation can be decomposed into a weighted summation of single-gene perturbations. Although we would expect that this combination of perturbations would induce a PAB/RVF-like transcriptomic effect, we can make no guarantee that this represents a unique solution and, more relevantly, that it represents the sequence of gene expression events that underly RVF. Nevertheless, we find it of interest that we were able to achieve such good fit performance and recommend further investigation into such inverse problem approaches.

The non-uniqueness of our solution merits further discussion. Our permutation analysis demonstrated that while the specific gene-weight pairings learned from the murine PAB data significantly outperform shuffled assignments in stratifying human RVF severity, randomly selected 41-gene perturbation sets with refit weights did not perform significantly worse at cross-species generalization (p = 0.109). This finding suggests that the transcriptional architecture of right ventricular failure may be sufficiently low-dimensional that multiple distinct perturbation combinations can approximate the disease transcriptome with comparable fidelity. Biologically, this is consistent with the notion of regulatory degeneracy, the principle that structurally distinct genetic perturbation sets can converge onto equivalent functional states through shared downstream effector pathways. Indeed, our observation that mitochondrial transcriptome dysregulation is a common convergent response to diverse perturbations provides a mechanistic basis for this degeneracy: if many different genetic insults ultimately funnel through a limited number of shared stress-response programs, then many different weighted combinations of those insults could plausibly reconstruct the integrated disease phenotype. From a practical standpoint, this degeneracy has important implications: it suggests that while the inverse problem may be solvable in the sense of achieving good transcriptomic fit, the specific solution should be interpreted as one of potentially many equivalent perturbation sets rather than as a unique mechanistic decomposition of the disease process.

We also note a tension between the assumptions underlying our reconstruction framework and our empirical findings on combinatorial genetic interactions. Our reconstruction model assumes strict additivity, specifically that the transcriptional effect of multiple simultaneous perturbations equals the weighted sum of individual effects. However, our analysis of second-order interactions demonstrates that combinatorial effects in cardiomyocytes are predominantly sub-linear, with the combined effect of two perturbations consistently falling below the sum of their individual effects. This systematic sub-linearity suggests that our linear model may overestimate the magnitude of the perturbation weights required to fit the disease transcriptome, or alternatively, that the model may require more perturbations than would be biologically necessary to achieve the observed phenotype. Future work incorporating non-linear interaction terms into the reconstruction framework (e.g. through interaction-aware regression or neural network-based approaches such as those proposed by Gonzalez et al.^23^) may improve both fit performance and the biological interpretability of the inferred perturbation sets.

Perhaps the most notable finding of this analysis is that perturbation weights derived exclusively from murine *in vivo* data, without any human training data, were able to stratify human RV samples by disease severity (Spearman ρ = 0.71, Kruskal-Wallis p = 0.016). This cross-species generalization implies that the transcriptional programs governing the transition from compensated pressure overload to overt right ventricular failure are at least partially conserved between mouse and human. While the relatively small size of the human validation cohort warrants caution in over-interpreting the magnitude of this effect, the fact that the observed stratification was significantly better than that achieved by permuted gene–weight assignments (p < 0.001) provides evidence that the specific perturbation signatures identified in our screen capture biologically meaningful transcriptional variation that generalizes across species. We emphasize, however, that transcriptomic conservation does not necessarily imply conservation of the upstream causal mechanisms, and that the perturbation weights should be interpreted as descriptive features of shared transcriptional covariation rather than as evidence that the same genetic perturbations drive RVF in both species.

From a translational perspective, the ability to decompose a complex disease transcriptome into a weighted combination of single-gene perturbation effects has potential implications for therapeutic target prioritization. If the perturbation weights learned by the model reflect, even approximately, the relative contribution of each gene’s dysregulation to the overall disease state, then genes with the largest absolute weights represent candidates whose correction might disproportionately restore the healthy transcriptome. In the context of RVF, this framework could in principle be used to predict which combinations of partial genetic corrections (achievable, for example, via AAV-delivered gene therapies or antisense oligonucleotides) would be most effective at reversing the disease transcriptional signature. Experimental validation of these predictions, for example by co-perturbing the top-weighted genes identified by our model and measuring whether the resulting transcriptome resembles the PAB/RVF state, represents a critical next step. More broadly, as perturbation atlases grow in scope and resolution, inverse problem approaches such as the one described here may provide a systematic, data-driven framework for identifying combinatorial therapeutic strategies for complex diseases.

## ACKNOWLEDGMENTS

IAK acknowledges the fellowship support of the Paul and Daisy Soros Fellowship for New Americans and the NIH/NIMH (F30 MH122076). KL was supported by the American Heart Association (#24PRE1195620). ZA and JE were supported by the CHOP Frontier Program. ZA was supported by the NIH (DK107667, HL152446). JE was supported by the NIH (5K08HL159311).

## AUTHOR CONTRIBUTIONS

IAK and KL conceived the project and designed the experiments with input from ZA. ZA supervised this research and experimental design. JE provided the murine PAB and human RVF dataset. W.Zhao, W.Zhu, W.Zhou, J.Liang, J.Li, and YY assisted with mouse husbandry and experiment execution. All authors contributed to data analysis and manuscript preparation.

## DECLARATION OF INTERESTS

The authors declare no competing interests.

## SUPPLEMENTARY FIGURE TITLES AND LEGENDS

**Figure S1.**
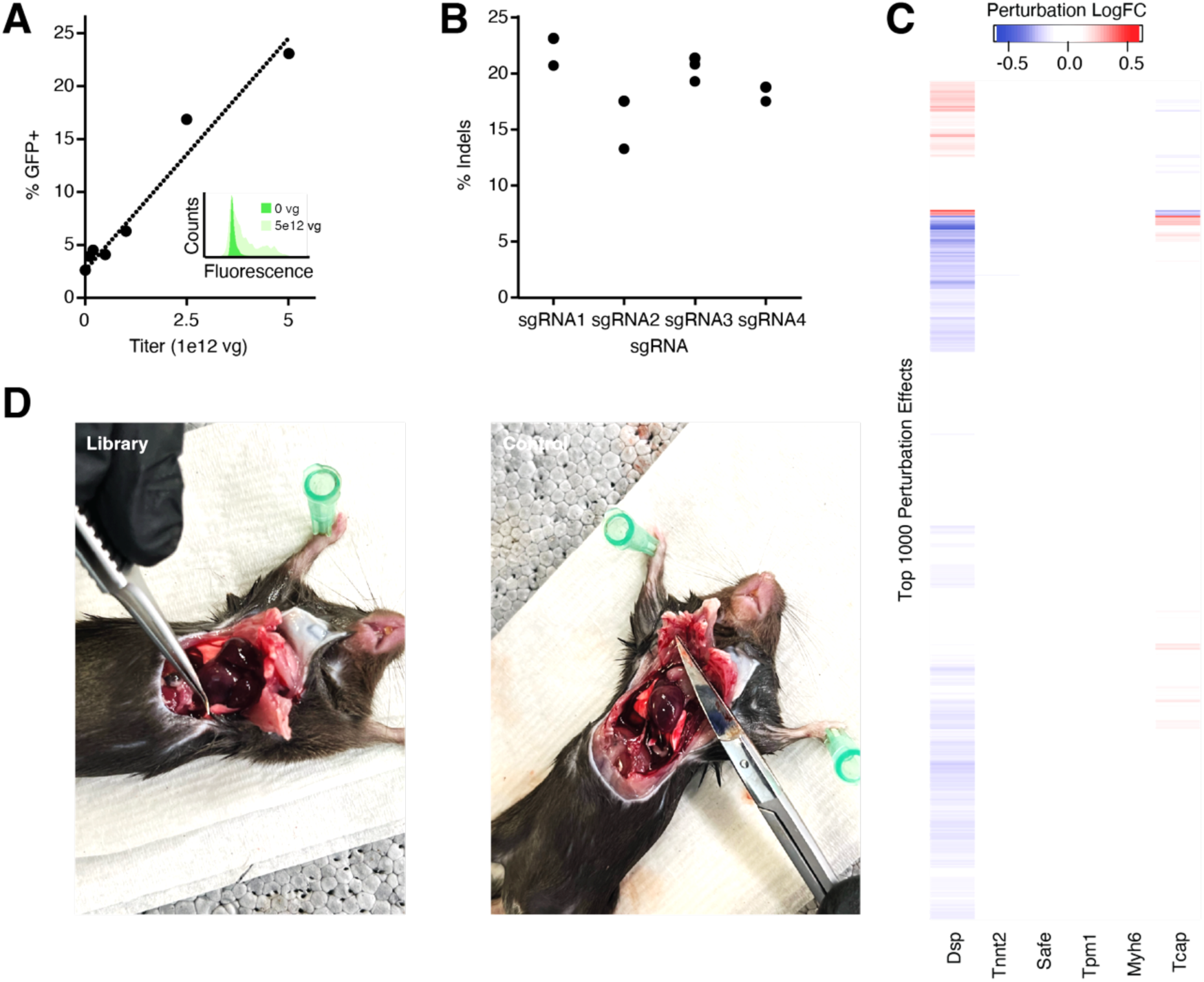
MyoAAV-mediated delivery allows for cardiac in vivo Perturb-seq, related to Figure 1. **(A)** In vivo titration of MyoAAV cardiac transduction efficiency shows a near linear relationship between viral titer and nuclear GFP positivity. **(B)** MyoAAV-mediated delivery of Pank1 sgRNA allows for efficient indel generation in the murine heart. **(C)** Heatmap of top 1000 perturbation effects after delivery of sgRNA for the corresponding gene. Insignificant effects (q < 0.2) are shaded white. **(D)** Delivery of sgRNA library at high MOI leads to cardiac chamber dilation.

**Figure S2.**
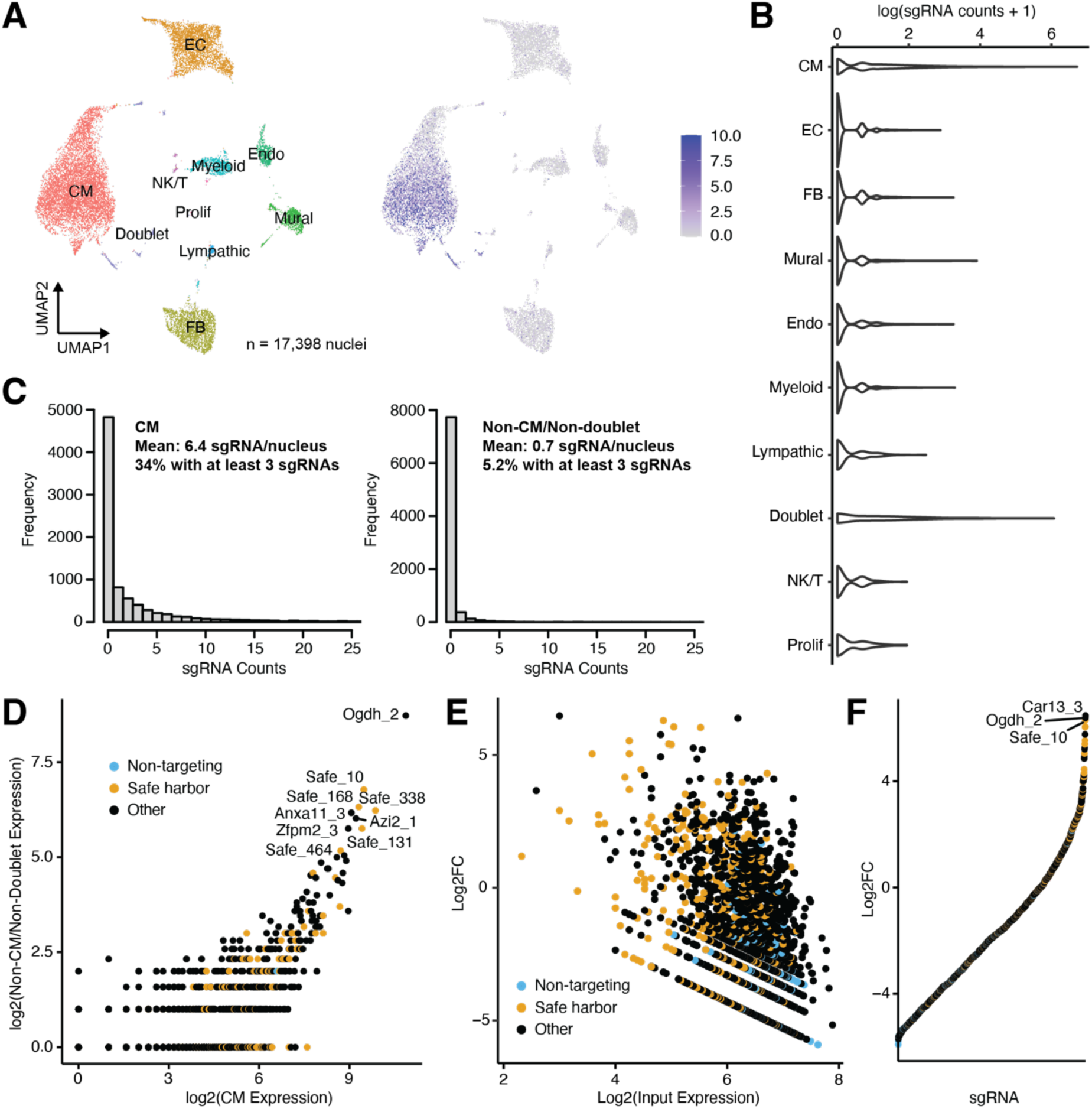
Technical characteristics of Perturb-seq screen, related to Figure 2. **(A)** UMAP showing isolated cell type identity (left) and colored by sgRNA count (right). **(B)** The majority of sgRNAs were assigned to cardiomyocyte or doublet nuclei. **(C)** Cardiomyocyte nuclei contained a mean of 6.4 sgRNA per nuclei, with ∼34% having at least 3 sgRNA (left). Non-cardiomyocyte/non-doublet nuclei contained a mean of 0.7 sgRNA per nuclei, with ∼5.2% having at least 3 sgRNA (right). **(D)** sgRNA assignment to non-cardiomyocytes correlated with the abundance of the same sgRNA in cardiomyocytes. **(E)** Observed fold-change in sgRNA abundance in cardiomyocytes versus in input library as a function of abundance in the input library. Note the trend for non-targeting guides to be poorly recovered from cardiomyocyte nuclei. **(F)** Ranked observed fold-change in sgRNA abundance in cardiomyocytes versus in input library.

**Figure S3.**
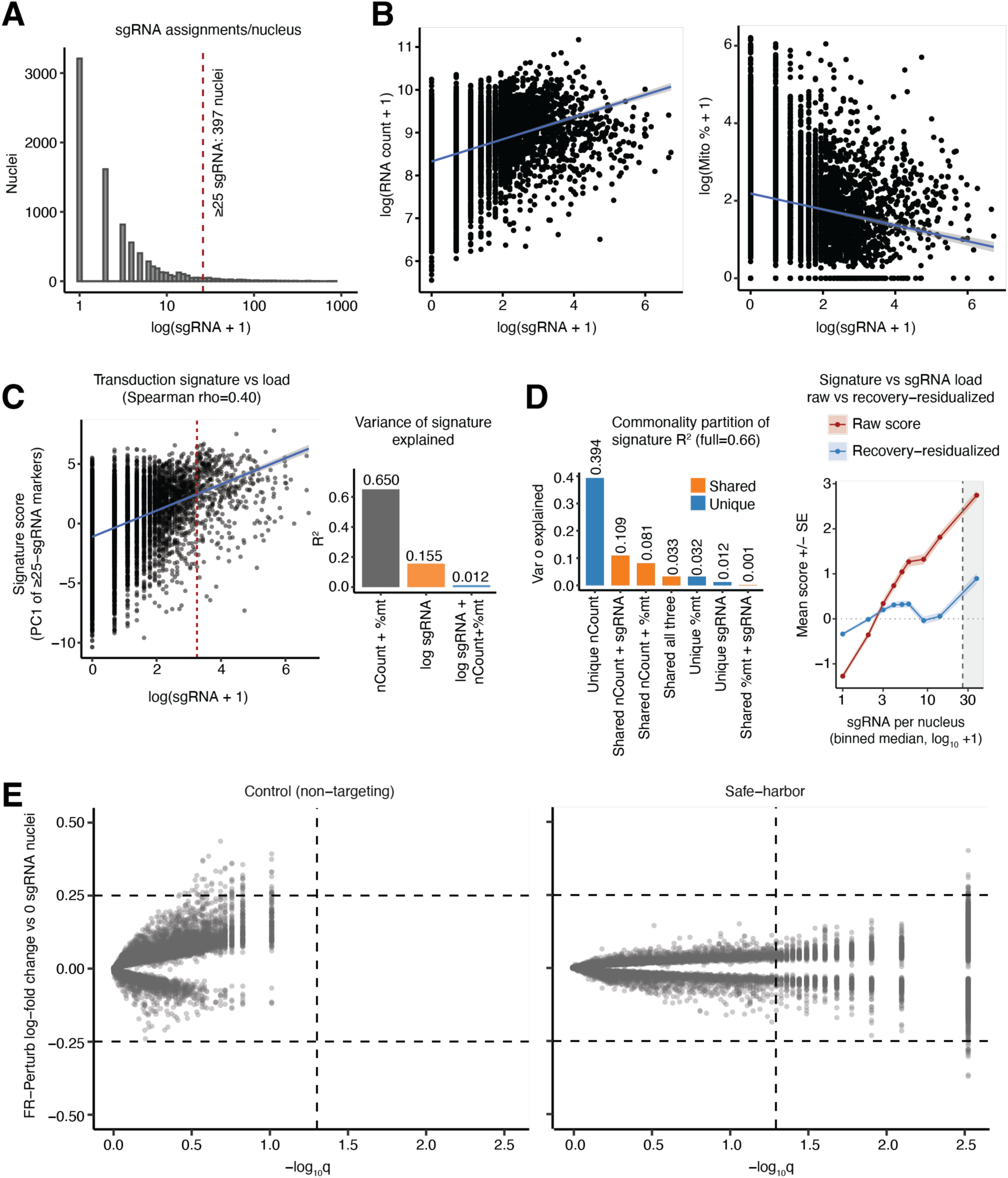
Apparent transduction-load transcriptional signature is overwhelmingly technical, and perturbation-effect estimation is robust to the control/normalization choice, related to Figure 2. **(A)** Distribution of assigned sgRNAs per cardiomyocyte nucleus (log_10_ scale); dashed line marks the ≥25-sgRNA “super-transducer” threshold (max 828 sgRNAs per nucleus). **(B)** Across cardiomyocyte nuclei, apparent sgRNA load is positively coupled to transcript recovery and inversely to mitochondrial content (Spearman ρ = +0.38 vs log nCount; ρ = −0.23 vs log %mito), i.e. guide and transcript capture covary. **(C)** A transduction-signature score (first principal component of the 406 genes upregulated in ≥25-sgRNA cardiomyocyte nuclei; **left**) is largely technical: total UMI count and mitochondrial fraction explain R² = 0.65 of its variance, whereas log sgRNA load adds only +1.2% beyond them (added R² = 0.012; **right**). **(D)** Commonality (variance) partition of the signature score (regressed on nCount, %mito and log sgRNA; R²_full = 0.66; **left**). Of sgRNA load’s gross association (R² ≈ 0.155) only 0.012 is unique to sgRNA whereas 0.109 is shared with recovery (nCount), i.e. its apparent effect is overwhelmingly collinear with transcript recovery. Mean signature score across binned sgRNA load, raw versus recovery-residualized (residuals of score ∼ nCount + %mito; **right**). Linear recovery adjustment removes ∼70–80% of the apparent dose-response (raw range ≈ 4.0 → residual ≈ 1.2; the residualized curve is flat across most of the sgRNA transduction range. The residual signal that remains is confined to the shaded marker-defining ≥25-sgRNA bin, where the relationship is circular (the 406 markers were defined from those nuclei). **(E)** FR-Perturb effects of the non-targeting “Control” and “Safe-harbor” perturbation columns (vs the 0 sgRNA reference) are near-null (0 and 30 genes at |LFC| > 0.25 and q < 0.05; median |LFC| ≈ 0.04), demonstrating that estimated perturbation effects are not an artifact of the control/normalization choice.

**Figure S4.**
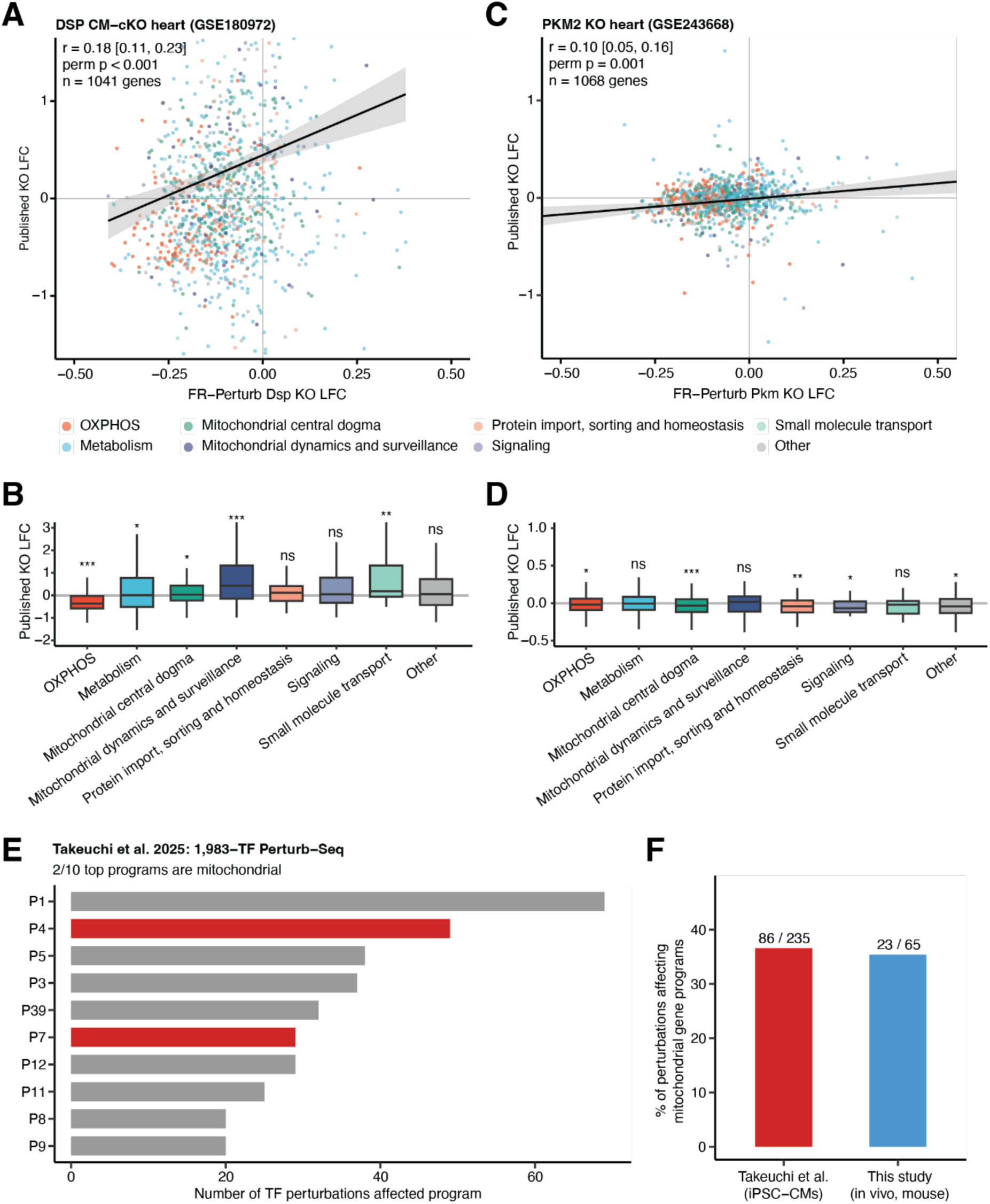
Mitochondrial transcriptome dysregulation is a conserved response to cardiac perturbation across independent screens, related to Figure 3. **(A)** Correlation between FR-Perturb log-fold changes (this study) and published bulk RNA-seq log-fold changes from DSP cardiomyocyte-specific conditional KO hearts (GSE180972). Each point represents one MitoCarta 3.0 gene, colored by mitochondrial functional category. Pearson r with bootstrap 95% CI and permutation p-value are shown. LFCs estimated using DESeq2. **(B)** Distribution of published DSP KO log-fold changes across MitoCarta 3.0 functional categories. Boxes show median ± IQR; asterisks indicate Wilcoxon signed-rank test against 0 (p < 0.05, ** p < 0.01, *** p < 0.001; minimum 5 genes per category). **(C-D)** As in (A-B) for PKM2 cardiac KO mice (GSE243668; PKM2^fl/fl^ versus KO at baseline). **(E)** The 10 gene programs most commonly disrupted across 1,983 TF perturbations in the Takeuchi et al. 2025 iPSC-cardiomyocyte Perturb-seq dataset, ranked by the number of TF knockdowns affecting each program. Programs with ≥10% overlap with MitoCarta 3.0 genes (top 300 program members) are highlighted in red. **(F)** Fraction of TF perturbations affecting at least one mitochondrial gene program in the Takeuchi et al. 2025 iPSC-CM screen (human, *in vitro*) versus this study (mouse, *in vivo*). Mitochondrial programs defined as described in (E); perturbations in this study classified using FR-Perturb q < 0.2.

**Figure S5.**
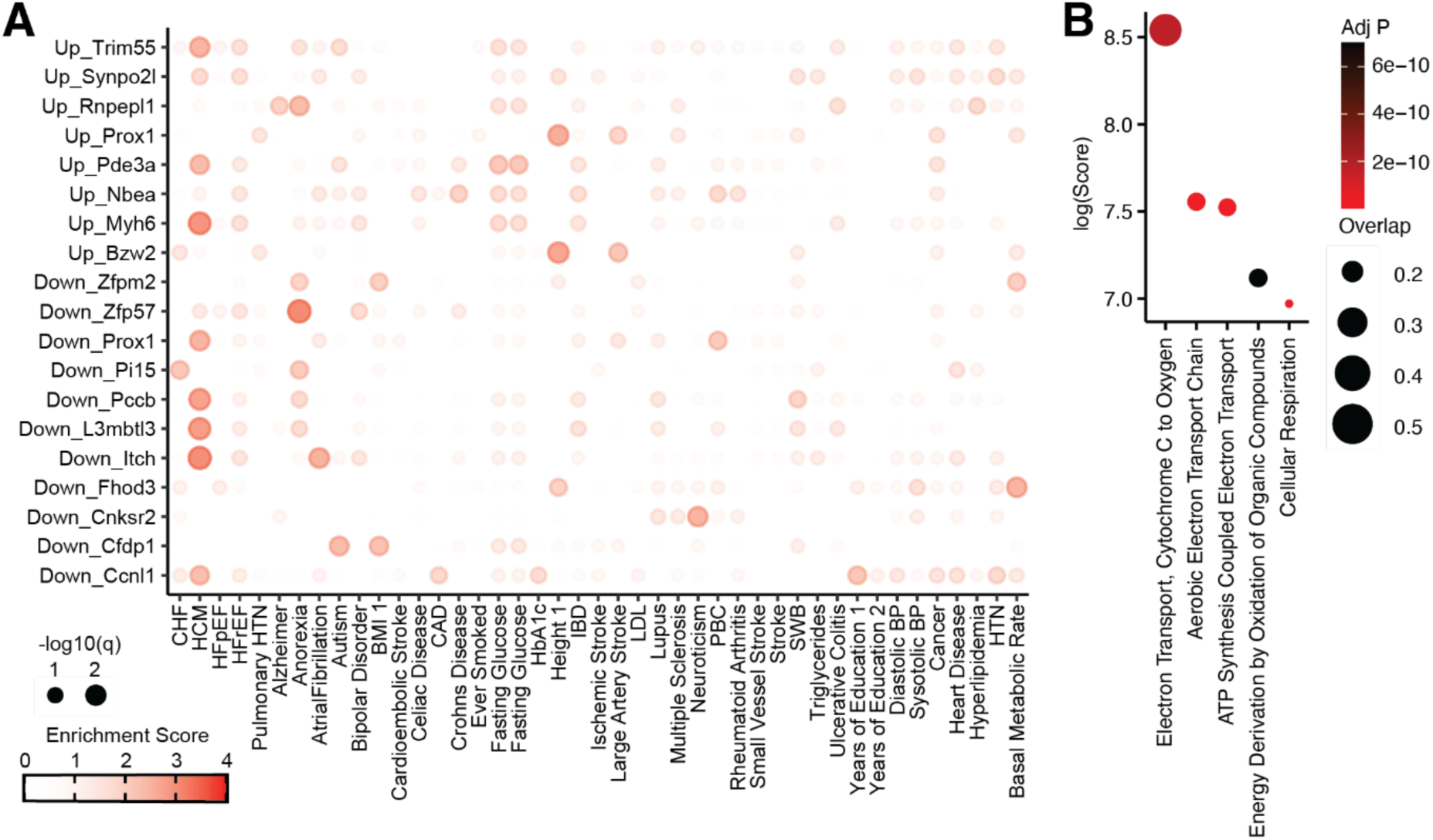
Integration of Perturb-seq with GWAS highlights the role of mitochondrial dysfunction in hypertrophic cardiomyopathy, related to Figure 3. **(A)** Sc-linker computed heritability enrichment scores of perturbation signatures (rows) across human traits (columns). Signatures are defined by a weighted collection of altered downstream genes for each perturbation. “Up” indicates that the corresponding signature is the set of genes that are upregulated by perturbation while “down” represents the opposite. Only perturbations/traits with at least one entry with q < 0.5 are shown. **(B)** Gene ontology enrichment of the union of the top 50 heaviest weighted downstream genes making up the signatures of perturbations with q < 0.2 for hypertrophy cardiomyopathy.

**Figure S6.**
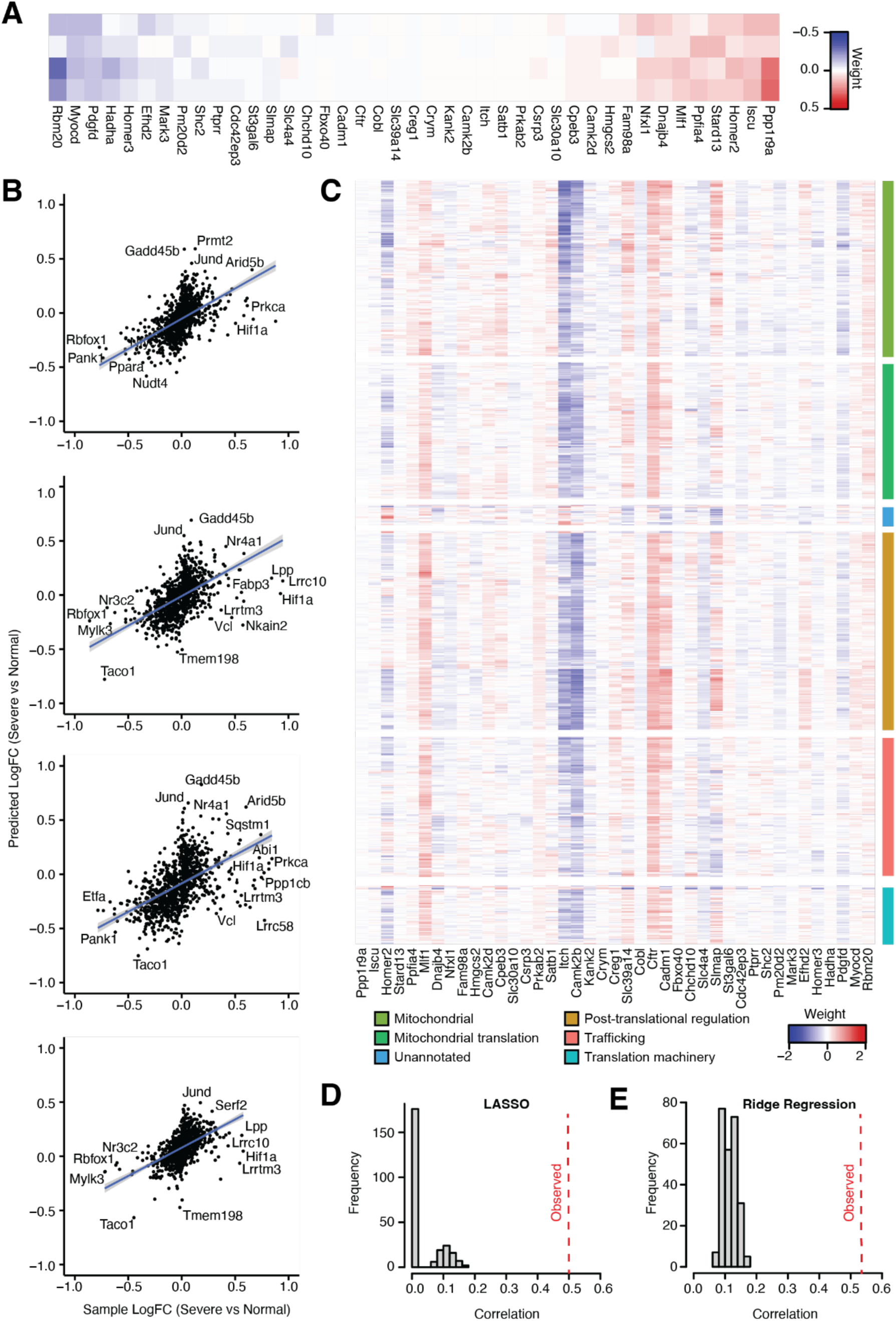
Reconstruction of the murine RVF transcriptome using gene perturbation signatures, related to Figure 4. **(A)** Heatmap showing weights of LASSO (alpha = 1) minimized perturbation-set which replicates the transcriptional signature of murine RVF. **(B)** Dot plots showing measured versus fit fold changes between healthy and failing murine RVs across 4 samples. **(C)** Heatmap showing effects of selected perturbations on downstream genes. The observed fit is significantly better, as measured by linear correlation between predicted vs measured fold change, than what would be observed by random chance, as calculated by randomly permuting downstream genes and re-fitting data with either LASSO **(D)** or ridge regression **(E)**.

## SUPPLEMENTARY DATA TITLES AND LEGENDS

**Data S1. Pilot sgRNA library, related to Figure 1**.

**Data S2. Full screen sgRNA library (585 genes), related to Figure 2**.

**Data S3. Gene modules, related to Figure 2**.

**Data S4. Gene Ontology modules, related to Figure 4**.

## METHODS

## RESOURCE AVAILABILITY

### Lead contact

Further information and requests for resources and reagents should be directed to and will be fulfilled by the lead contact, Zoltan Arany.

### Materials availability

Mouse lines and plasmids are available upon request from the lead contact.

### Data and code availability

- Code statement: The sequencing analysis pipeline, as well as code to generate paper figures, is available at Github: https://github.com/ikuznet1/Cardiac_InVivo_Perturb and Zenodo DOI: 10.5281/zenodo.20495835.
- Any additional information required to reanalyze the data reported in this paper is available from the lead contact upon request.

## EXPERIMENTAL MODEL AND SUBJECT DETAILS

### Mouse models

All animal studies were approved by the University of Pennsylvania Institutional Animal Care and Use Committee (protocols 805255 and 805309). Mice were housed on a normal light-dark cycle (light on from 7 am to 7 pm) with constant access to normal chow (Lab diet, 5010) and water, unless otherwise specified. Experiments were initiated on mice of 6-10 weeks (typically 7-8 weeks) of age, unless otherwise stated. All experiments were performed with littermate controls. All mouse procedures were approved by the University of Pennsylvania Animal Care and Use Committee. The mouse studies described here involved LSL-Cas9 (JAX strain #024857) mice. All mice were male. Nuclei from all four animals were pooled and distributed across all 10x lanes prior to droplet generation; animal identity was not multiplexed and is therefore not recoverable at the single-nucleus level. Consequently, animal is not confounded with lane or sequencing batch, but between-animal variability cannot be estimated. The primary endpoint is a quantitative, sequencing-based computational readout (FR-Perturb) and is operator-independent; blinding is not applicable to the primary analysis. Histological and gross-morphological assessments were performed not blinded to condition.

## METHOD DETAILS

### Single-nuclei sample preparation

Samples were prepared as described previously. ^27–29^ Briefly, frozen cardiac LV or RV were sectioned to 100 μm (Leica CM1950 cryostat) and then transferred into a 2 mL Dounce homogenizer containing 1 ml of chilled lysis buffer (10 mM Tris-HCl, pH 8.0, 250 mM sucrose, 25 mM KCl, 3 mM MgCl_2_, 1 μM DTT, DAPI (1 μg/mL), and 0.05% IGEPAL-630 in nuclease-free water). Samples were homogenized with ten passes of a course pestle, then ten passes of a fine pestle, and then incubated on ice for 15 min. Large debris was removed by centrifugation at 40g for 1 minute (Beckman Coulter Allegra X-15R swinging bucket centrifuge) before sequentially being filtered through a 50-μm and then 10-μm filter (puriSelect Life Science) into a 50-ml conical tube. The filters were washed with 6 mL of Wash Buffer (0.01% BSA, 5mM MgCl_2_, PBS) and the resultant suspension was pelleted at 550g for 5 minutes at 4 °C. The pelleted nuclei were resuspended in 150 μL of cold Flow Buffer (1% BSA, 0.4 U/μ murine RNAse inhibitor (New England Biolabs), PBS). GFP+ nuclei were enriched via fluorescence activated cell sorting (FACS). Nuclei were flowed at the lowest flow rate on a BDFACSARIA III with a 100-μm nozzle. DAPI+/GFP+ singlets were collected. Nuclei were recovered by spinning at 500x g for 5 minutes and then resuspended in Nuclei Resuspension Buffer before quantification and dilution to 1,000 nuclei μl^−1^.

### Power analysis

Yao et al. reported an ∼11.25 increase in efficiency with guide-pooling as opposed to conventional Perturb-Seq.^8^ Using a library of similar complexity as ours, they were able to power a conventional Perturb-Seq screen with 60,000-90,000 droplets, thereby suggesting similar power could be achieved in a compressed Perturb-Seq screen with 5,300-8,000 droplets. This efficiency gain was achieved with a mean of 2.50 sgRNAs per droplet and with 35% of droplets containing 3 or more sgRNAs.

### ECCITE-Seq & 10X Chromium Single Cell Gene Expression

Nuclei were loaded onto the 10x Genomics Chromium microfluidic platform (Single cell 5′ solution, v2) for an estimated recovery of 5,000 cells per device. Processing of libraries was performed according to manufacturer’s instructions with some notable exceptions as described previously. Specifically, prior to the reverse transcription step an additional primer was spiked into the cocktail to generate sgRNA cDNA. During the SPRI selection steps the libraries were separated by size into a cDNA library and a sgRNA library. sgRNA libraries were amplified and barcoded as previously described.

### Sequencing & alignment

cDNA libraries were multiplexed and sequenced on a NovaSeq 6000 S4 chip to a goal of ∼75000-100000 reads per nuclei. Raw base call files for each sample were de-multiplexed and converted to FASTQ files using the 10x Genomics toolkit CellRanger 9.0.1 (cellranger mkfastq). FASTQ files were then uploaded to the 10x Genomic Cloud where Cellranger Count 7.1.0 was run on each sample to map them to the Mouse (GRCm39) 2024-A transcriptome with the include-introns flag set to true. Resultant raw feature matrices were then used for downstream analysis. sgRNA libraries were similarly sequenced on a NovaSeq 6000 S4 chip. Resultant FASTQ files were then analyzed via *CITE-seq-Count*.^30^

### Filtering and quality control

Cell-type assignment was conducted in *Seurat*.^31^ Nuclei with >5% mitochondrial transcripts, with <200 transcripts, or with >100,000 transcripts were discarded. Transformation and normalization was performed using SCTransform, with mitochondrial read percentage regressed out. 50 principal components were utilized for downstream analysis. For doublet removal, we elected to utilize a semi-supervised approach, as has been done previously,^32^ which was based upon identifying nuclei which simultaneously had gene expression signature consistent with two (or more) cell populations. Practically, this was done by iteratively clustering nuclei at high resolution, generating z-score expression profiles for each expected cell population, and visualizing the nuclei clusters colored by z-score using UMAP embedding. Clusters demonstrated high expression profiles for multiple cell types were deleted and then entire above process was repeated until no further clusters were removed (six rounds).

### sgRNA-to-nuclei assignment

The output of CITE-seq-Count yielded a matrix with rows corresponding to nuclear barcodes, columns corresponding to sgRNAs, and values corresponding to the UMI count for a specific sgRNA in that specific nucleus. Due to ambient RNA contamination, a certain degree of incorrect assignment of sgRNA to nucleus is to be expected. We tested multiple different strategies of assigning sgRNAs to nuclei via *crispat*.^33^ Using the generated sgRNA assignments we then ran multiple iterations of FR-Perturb with cross-validation and identified the sgRNA assignment strategy that delivered the best performance. Empirically, this turned out to be utilizing an absolute UMI cutoff of at least 3 to consider an sgRNA as being present in a droplet.

### FR-Perturb

FR-Perturb was run with the control perturbation as 0 sgRNA droplets. The percentage of mitochondrial reads, RNA count, and cardiac chamber of origin were regressed out. The “--guide-pooled” flag was set to true.

### Covariate selection for FR-Perturb effect estimation

Perturbation effects were estimated with FR-Perturb including per-cell total UMI count (nCount_RNA), mitochondrial transcript fraction (percent.mt), and sample of origin as technical covariates; per-cell sgRNA count was deliberately not included as a covariate. This choice was guided by an explicit decomposition of the transcriptional variance associated with apparent transduction load (**Figure S3C–D**). We defined a transduction-signature score as the first principal component of the 406 genes upregulated in heavily transduced cardiomyocyte nuclei (≥25 assigned sgRNAs versus the remainder; log_2_ fold-change > 0, Benjamini–Hochberg adjusted P < 0.05). nCount_RNA and percent.mt together explained 65.0% of the variance in this score (**Figure S3C**); a commonality analysis (**Figure S3D**) attributed 39.4% of score variance uniquely to nCount_RNA and 3.2% uniquely to percent.mt, indicating that the apparent transduction-correlated expression program is overwhelmingly a recovery/quality axis, and these variables were therefore retained. Apparent sgRNA load explained little additional variance: log sgRNA count contributed only 1.2% beyond nCount_RNA and percent.mt (added R² = 0.012; **Figure S3C**), and commonality analysis showed that ∼70% of its gross association with the signature (0.109 of 0.155) was shared with nCount_RNA rather than unique (**Figure S3D**), consistent with the expected positive coupling between guide and transcript recovery (Spearman ρ = 0.38; **Figure S3B**). sgRNA count was thus uninformative as a technical covariate. More fundamentally, per-cell sgRNA count is a proxy for the perturbation exposure itself so adjusting for it would constitute over-adjustment for the treatment whose effect is being estimated and would remove biological signal of interest. We therefore modeled perturbation identity (the FR-Perturb design) while conditioning only on the recovery covariates (nCount_RNA, percent.mt) and batch.

### Second-order effects

We calculated intra-modular and inter-modular second-order effects as previously described.^8^ Briefly, to calculate intra-modular effects, we first grouped perturbations into 146 (potentially overlapping) GO annotation modules. Separately, we grouped downstream perturbed genes into gene programs. For each module/program pair we defined 3 sets: 1) control set – cells containing only non-targeting sgRNAs, safe harbor locus targeting sgRNAs, or sgRNAs without significant effects on the given program; 2) first-order effect set – cells containing exactly one sgRNA from the given module with all other sgRNAs in that cell contained with the control set; 3) second-order effect set – cells containing exactly two sgRNAs from the given module with all other sgRNAs in that cell contained with the control set. We then computed a mean expression value for each program in each set (denoted *μ*_0_, *μ*_1_ and *μ*_1,1_) as the average log(TP10K + 1) expression of all genes in the given program. We regressed read count per cell and percent mitochondrial reads and standardized the log(TP10K + 1) expression to mean 0 and variance 1 prior to running this calculation. We then define first-order effects as *β*_1_ = *μ*_1_ – *μ*_0_ and second-order effects as *β*_1,1_ = *μ*_1,1_ – 2*β*_1_ – *μ*_0_. Inter-modular effects were computed similarly, with the exception that we grouped our GO modules into 8 super-modules via Leiden clustering. For each triplet (i.e. a super-module pair and a gene program) we again defined 4 sets: 1) control set – cells containing only non-targeting sgRNAs, safe harbor locus targeting sgRNAs, or sgRNAs without significant effects on the given program; 2) first-order effect set A – cells containing exactly one sgRNA from one of the super-modules (but not both) with all other sgRNAs in that cell contained with the control set; 3) first-order effect set B – cells containing exactly one sgRNA from the other super-module (but not both) with all other sgRNAs in that cell contained with the control set; 4) second-order effect set – cells containing exactly one sgRNA from each super-module with all other sgRNAs in that cell contained with the control set. We calculate the mean expression of each set as before (denoted *μ*_0_, *μ*_1_, *μ*_2_, and *μ*_1,2_). We then define first-order effects as *β*_1_ = *μ*_1_ – *μ*_0_, *β*_2_ = *μ*_2_ – *μ*_0_, and second-order effects as *β*_2,1_ = *μ*_1,2_ – *β*_1_ – *β*_2_ – *μ*_0_. We calculate P values by permuting set membership labels of cells and recomputing *μ* and *β*. We calculate standard errors by bootstrapping, resampling cells from each set.

### GWAS integration

We initially constructed two programs for each perturbed gene which either reflected a weighted set of upregulated or downregulated downstream genes. Weights were calculated via linearly scaling log2 fold-changes to a maximum value of 1. We retained programs with at least 50 downstream genes, yielding 62 upregulated programs and 61 downregulated programs. We then assumed that if a perturbed gene was important for a given trait, then disease heritability should be enriched near its downstream perturbed genes. We then used sc-linker, as previously described, to estimate disease heritability enrichment scores for each gene program.^8,17^

### Reconstructing the murine RVF transcriptome

We acquired snRNA-seq data from mice which had undergone sham (n = 3) vs pulmonary artery banding (PAB; n = 3 moderate RVF and n = 4 severe RVF; a model of RV pressure overload). We then fit the log(TP10K + 1) normalized transcriptomic shift *y* between PAB and sham cardiomyocytes as a weighted summation of perturbations. We restricted our analysis to gene which were differentially expressed between the sham and PAB groups. Initially, rather than using single-gene perturbation effects, we instead defined GO-based perturbation modules (using our previously combined GO modules) as the average perturbation effect of all module member genes. We then used elastic net regression to fit the weights. We conducted model fitting in a lower-dimensional subspace defined by the initial 5 principal components of the matrix generated by concatenating GO-module perturbation effects. We then sought to find a minimal set of weighted single-gene perturbation effects which could effectively fit the PAB transcriptome. We restricted our analysis to the perturbed genes which were present in the GO-based perturbation modules with non-zero weights in our prior analysis. We then fit *y* as a weighted combination of single-gene perturbation responses, using LASSO regression to optimize weights while enforcing sparsity. We calculated the null distribution by permuting gene labels for measured perturbation effects.

### Histological analysis

Cardiac tissues were fixed in 4% PFA for at least 16 hours and then dehydrated into 100% ethanol. Tissues were subsequently embedded in paraffin by the CVI Histology Core and sectioned. H&E staining was performed with CAT Hematoxylin (Biocare Medical, CATHE-M) and Edgar Degas Eosin (Biocare Medical, THE-MM). GFP was stained with ab6673.

### Enrichment Analysis

Enrichment analysis was conducted with *EnrichR*,^34^ relying heavily on the ChEA^35^ and Gene Ontology^36^ datasets.

## QUANTIFICATION AND STATISTICAL ANALYSIS

Details of specific statistical analyses for each section, sample sizes, and statistical tests used are given in the methods and in the corresponding figure legends. Notably, all analysis was done in R using primarily custom written scripts and the Seurat package, which are available with deposited data.

## Notes

https://zenodo.org/records/20495835

https://github.com/ikuznet1/Cardiac_InVivo_Perturb

### Competing Interest Statement

The authors have declared no competing interest.

